# ARS2-directed alternative splicing mediates CD28 driven T cell glycolysis and effector function

**DOI:** 10.1101/2021.05.07.442963

**Authors:** G. Aaron Holling, Anand P. Sharda, Mackenzie M. Honikel, Caitlin M. James, Shivana M. Lightman, Guanxi Qiao, Kelly L. Singel, Tiffany R. Emmons, Thejaswini Giridharan, Shengqi Hou, Andrew M. Intlekofer, Richard M. Higashi, Teresa W. M. Fan, Andrew N. Lane, Kevin H. Eng, Brahm H. Segal, Elizabeth A. Repasky, Kelvin P. Lee, Scott H. Olejniczak

## Abstract

CD8 T cell activation prompts extensive transcriptome remodeling underlying effector differentiation and function. Regulation of transcriptome composition by the mitogen-inducible nuclear cap-binding complex (CBC) adaptor protein ARS2 has critical cell type-specific consequences, including thymic T cell survival. Here we show that ARS2 was upregulated by CD28 during activation of peripheral T cells, was essential for anti-tumor immunity, and facilitated T cell activation-induced alternative splicing. The novel splicing function of ARS2 was mediated at least in part by recruitment of splicing factors to nascent transcripts including the M2 isoform of pyruvate kinase (*Pkm2*), a key determinant of CD8 T cell effector properties. Notably, ARS2-directed *Pkm2* splicing occurred days after stimulation of PI3K-indepdendent CD28 signaling and increased glycolysis beyond levels determined by PI3K signaling during T cell priming. Thus, ARS2-directed *Pkm2* splicing represents a mechanism by which CD28 drives glycolytic metabolism, allowing for optimal effector cytokine production and T cell anti-tumor immunity.

## Introduction

Anti-tumor CD8 T cell responses rely on coordinated regulation of transcriptome output, enabling growth, proliferation, and acquisition of effector properties^1^. Such bioenergetically demanding processes are enabled by multi-phase metabolic reprogramming that occurs following T cell activation^2^. Glycolysis is induced within 24 hours of activation through PI3K-dependent upregulation of glucose transporters^3^. Continued glycolytic flux drives production of the key effector cytokine interferon gamma (IFNγ) by limiting binding of the glycolysis enzyme GAPDH to *Ifng* mRNA^4^. Several recent studies suggest that another glycolysis enzyme, PKM2 also has a “moonlighting” function in T cells; PKM2 translocates to the nuclei of activated CD4 T cells where it regulates transcription and thus affects Th17 differentiation^5, 6, 7^. A direct transcriptional regulatory function of PKM2 has also been reported in tumor cells, although this non-metabolic function remains controversial^8^. In tumor cells, PKM2 expression is regulated by mutually exclusive alternative splicing of exons 9 (*Pkm1*) and 10 (*Pkm2*) as the result of oncogene-driven changes in expression of splicing factors including SRSF3, hnRNPA1/A2, and PTBP1^8^. The process of alternative splicing has gained some appreciation as a mechanism that regulates T cell responses^9^, yet whether *Pkm* splicing plays a role in T cells was unclear.

RNA polymerase II transcripts are modified with a 7-methyguanosine cap at the 5’-end that recruits the nuclear cap-binding complex (CBC). The CBC, composed of CBP80 and CBP20, directs virtually all aspects of RNA processing including splicing, 3’-end processing, nuclear export, and translation^10^. The CBC adaptor protein ARS2 serves as a scaffold between the CBC and RNA processing factors involved in RNA 3’ end processing and nuclear export, or RNA degradation^11, 12, 13^. ARS2 is an essential protein that plays context-specific roles in mammalian physiology, including maintenance of neural and hematopoietic stem cell identity ^14, 15^, retinal progenitor cell proliferation^16^, and thymocyte survival^15^. Unlike other CBC components, ARS2 expression fluctuates greatly in response to extracellular stimuli^17, 18, 19^, suggesting that regulation of ARS2 expression may be a means for cells to coordinate signaling cues with transcriptome output.

In the current study we set out to investigate the role of ARS2 in CD8 T cells. We found that ARS2 was induced by T cell activation in a CD28 co-stimulation dependent manner and was necessary for antigen-specific control of tumor growth in mouse models. Importantly, we report a novel function of ARS2 in regulating alternative splicing during T cell activation. Among ARS2 splicing targets, we identified PKM2 as a key regulator of CD8 T cell glycolytic reprogramming, IFNγ production, and anti-tumor immunity. Mechanistically, CBC-bound ARS2 recruited splicing factors SRSF3 and hnRNPA1 to *Pkm* pre-mRNA to promote *Pkm2* splicing. Finally, we found that ARS2-directed *Pkm2* splicing was regulated by the membrane distal CD28 PYAP intracellular signaling domain responsible for PI3K-independent CD28 signaling. Rescue of ARS2 or PKM2 in CD28 PYAP domain mutant T cells restored glycolysis and IFNγ production. Together, data identify a novel CD28-ARS2-PKM2 axis in CD8 T cells that drives glycolytic metabolism and effector cytokine production in support of anti-tumor immunity.

### The ARS2-bound CBC (CBCA) supports CD8 T cell anti-tumor immunity

Unlike in other cell types where ARS2 is the only regulated CBCA component, T cell activation resulted in upregulation of ARS2 (*Srrt*), CBP80 (*Ncbp1*), and CBP20 (*Ncbp2*) protein (Fig. 1a) and mRNA (Fig. 1b). Analysis of public datasets revealed CBCA upregulation was similar in CD4 and CD8 T cells (Extended Data Fig. 1a), unaffected by biological gender (Extended Data Fig. 1b), occurred following antigen stimulation of CD8 T cells (Extended Data Fig. 1c), and was enriched during early CD8 T cell effector differentiation (Supplementary Table 2 of Kakardov *et al*.^20^). To examine signaling requirements for CBCA upregulation T cells were stimulated with αCD3, αCD28, or both. αCD3 or αCD28 did not induce upregulation individually, while co-stimulation with αCD3 + αCD28 induced expression of *Srrt* and *Ncbp2* (Fig. 1c). Notably, *Srrt* upregulation required *de novo* transcription (Extended Data Fig. 1d) and occurred with greater magnitude than *Ncbp2* upregulation. These data are consistent with a role for CBCA upregulation in the cellular response to CD8 T cell activation.

**Figure 1:**
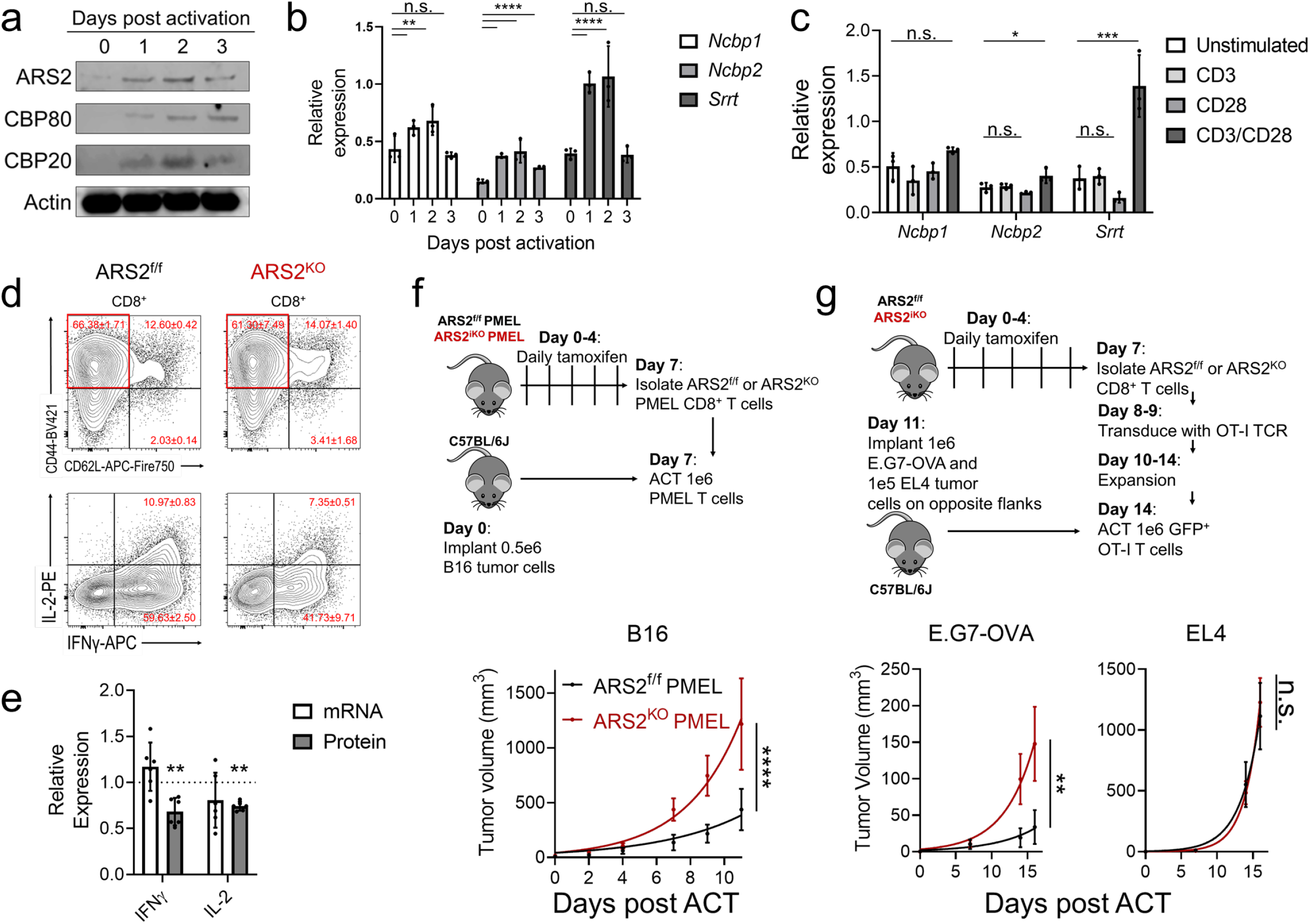
ARS2 supports CD8 T cell anti-tumor immunity. a) Western blots showing expression of CBCA components in human T cells stimulated with αCD3/αCD28 for indicated times. Representative of two healthy donors. b) Expression of transcripts coding CBCA components CBP80 (*Ncbp1*), CBP20 (*Ncbp2*), and ARS2 (*Srrt*) in C57BL/6J mouse T cells stimulated with αCD3/αCD28 + rIL-2 for indicated times. Graphs show mean qRT-PCR ΔCt value ± SD with *Tbp* as the endogenous control, dots represent independent mice. Unpaired t test, n.s.= not significant, **p < 0.01, ****p < 0.001. c) Expression of transcripts coding CBCA components in C57BL/6J mouse T cells stimulated with αCD3, αCD28, or αCD3 + αCD28 for 24 hours. Graphs show mean ± SD, dots represent independent mice. One-way ANOVA, n.s.= not significant, *p < 0.05, ***p < 0.001. d) Flow cytometry assessment of CD44 vs. CD62L expression on activated ARS2^fl/fl^ or ARS2^KO^ CD8 T cells (top) and IFNγ vs. IL-2 expression in effector T cells (CD8^+^CD44^+^CD62L^-^) following re-stimulation with PMA/ionomycin in the presence of brefeldin-A. Red text indicates mean % in each region ± SD of 4-6 mice/group. e) IFNγ and IL-2 protein (from **d**) and mRNA expression in ARS2^KO^ T cells relative to ARS2^fl/fl^ controls. Graphs show mean ± SD, dots represent individual mice. Unpaired t test, **p < 0.01. f) Growth of B16 melanomas in C57BL/6J mice following adoptive transfer of PMEL TCR transgenic CD8 T cells (ACT). Graphs show best fit lines to mean tumor volume ± SEM in mice receiving ARS2^f/f^ (black line, n = 6) or ARS2^KO^ (red line, n = 7) PMEL T cells. Two-way ANOVA, ****p < 0.0001. g) Growth of E.G7-OVA and EL4 tumors on opposite flanks of C57BL/6J mice following adoptive transfer of CD8 T cells engineered to express the OT-I TCR. Graphs show best fit lines to mean tumor volume ± SEM in mice receiving OT-I transduced ARS2^f/f^ (black line, n = 8) or ARS2^KO^ (red line, n = 7) T cells. Two-way ANOVA, n.s.=not significant, **p < 0.01.

To directly test CBCA in peripheral T cells we employed a tamoxifen inducible ARS2 knockout system (ARS2^iKO^) because constitutive ARS2 knockout in developing thymocytes resulted in apoptosis and failure of mature T cells to reach the periphery^15^. CBP80 and CBP20 inducible knockout mice do not currently exist. Tamoxifen induced *Srrt* deletion had no effect on the number or viability of mature splenic T cells (Extended Data Fig. 1e,f), upregulation of T cell activation markers CD69 and CD44 (Extended Data Fig. 1g), or activation-induced cell growth (Extended Data Fig. 1h), indicating ARS2 was not required for survival or activation of peripheral T cells. Further characterization revealed that ARS2^KO^ CD8 effector T cells (CD8^+^CD44^+^CD62L^-^) produced less IFNγ and IL-2 than controls (Fig. 1d and Extended Data Fig. 1i) despite equivalent expression of *Ifng* and *Il2* mRNAs (Fig. 1e).

Since effector cytokines are indicators of effector differentiation and are required for the cytotoxic activity of CD8 T cells, we examined the effect of ARS2 on CD8 T cell anti-tumor immunity. ARS2^iKO^ mice were bred to PMEL TCR transgenic mice to generate ARS2^iKO^ CD8 T cells that recognize the gp100_25-33_ antigen presented by H2-D^b^ on B16 melanoma cells^21^. Adoptively transferred PMEL TCR transgenic ARS2^KO^ CD8 T cells were unable to control growth of B16 melanomas to the same extent as ARS2 ^f/f^ controls (Fig. 1f). To replicate these findings in a clinical adoptive therapy model, ARS2^f/f^ or ARS2^KO^ CD8 T cells were isolated and transduced with the alpha and beta chains of the OT-I TCR^22^, thereby generating MHC class I-restricted OVA-specific CD8 T cells. Equal numbers of OT-I transduced CD8 T cells, determined by GFP co-expression (Extended Data Fig. 1j), were adoptively transferred into tumor-bearing mice with EL4 lymphoma and the OVA-expressing EL4 variant E.G7-OVA implanted on opposite flanks. Adoptively transferred OT-I transduced CD8 ARS2^KO^ T cells were unable to control growth of E.G7-OVA tumors as well as ARS2^fl/fl^ controls. EL4 tumors lacking OVA antigen expression grew at a similar rate in all mice (Fig. 1g). Together data indicate that induction of ARS2 expression supports CD8 T cell anti-tumor effector function.

### ARS2 regulates T cell activation-induced alternative splicing

To gain insight into how ARS2 affected T cell function we performed RNA-sequencing using WT and ARS2^KO^ T cells immediately following isolation (day 0) and following stimulation with αCD3/αCD28 microbeads + recombinant murine interleukin-2 (rIL-2) for 24 hours (day 1) or 72 hours (day 3). Surprisingly, the vast majority of differentially expressed genes (DEG) in activated T cells were unaffected by ARS2 deletion (Fig. 2a). Rather, PSI-Sigma^23^ analysis revealed that between one-quarter and one-third of alternative splicing events induced by T cell activation were disrupted in ARS2^KO^ T cells (Fig. 2a). Less common splicing events alternative 5’ and 3’ splice site utilization and mutually exclusive splicing were particularly sensitive to ARS2 deletion (Fig. 2b). These data indicate a novel splicing function of ARS2 during T cell activation, as ARS2 has not been shown to affect alternative splicing in immortalized cell lines despite identification of multiple splicing factors in ARS2 interactome studies^12, 13, 24^.

**Figure 2:**
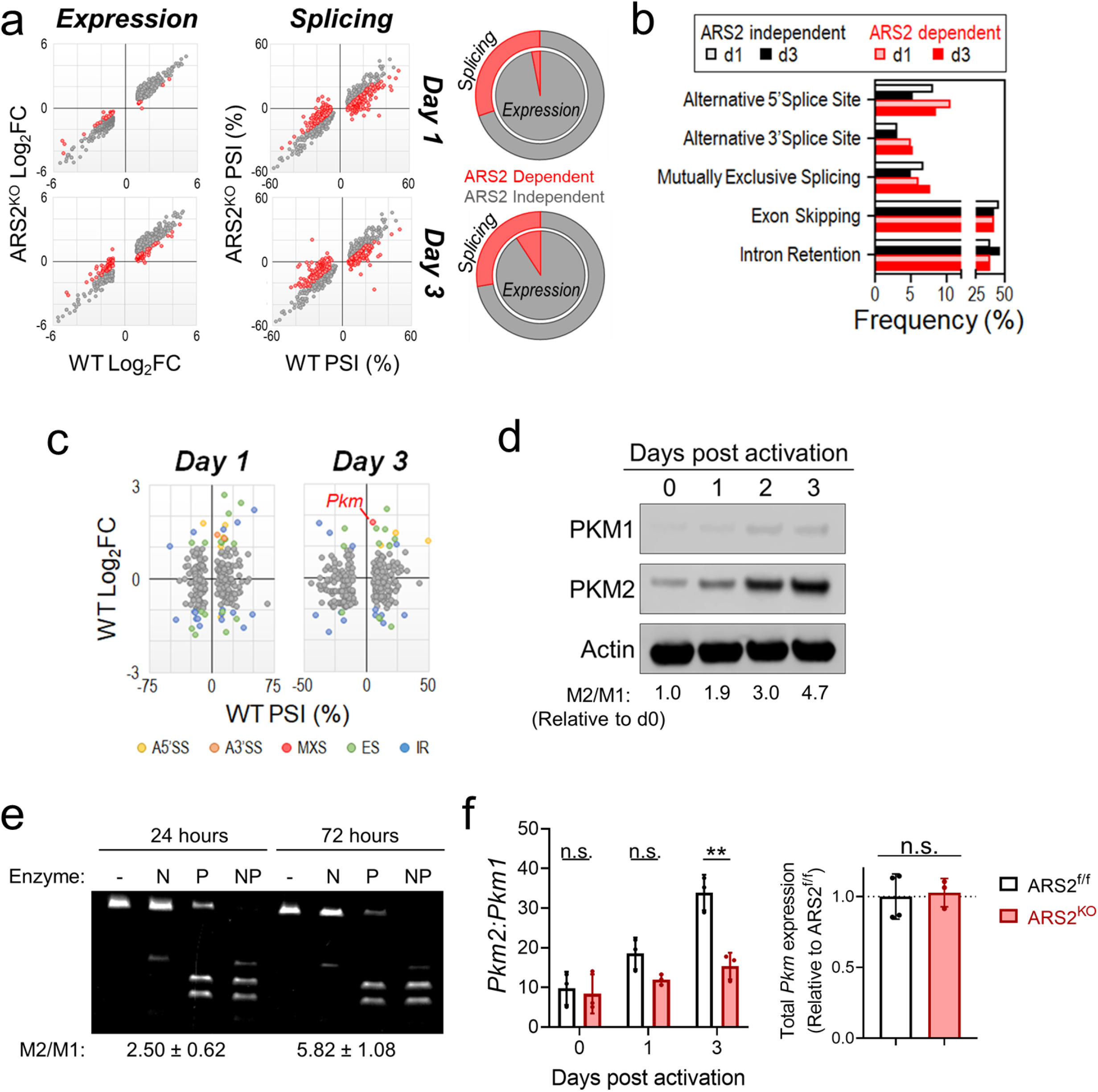
Regulation of alternative splicing by ARS2. a) Effect of ARS2 deletion on T cell activation-induced gene expression (right) or alternative splicing (left) at day 1 (top) and day 3 (bottom) following stimulation with αCD3/αCD28 + rIL-2. Red dots indicate significant differences between WT and ARS2^KO^ T cells. Pie charts depict the contribution of ARS2 to gene upregulation (inner pie) and alternative splicing (outer pie) induced by T cell activation. b) Prevalence of types of alternative splicing events induced by T cell activation and regulation by ARS2. c) Effect of T cell activation on expression (y-axis) vs. alternative splicing (x-axis) of genes whose alternative splicing was ARS2-dependent. Dots colored by type of splice event were induced more than two-fold by T cell activation. d) Western blots for PKM1 and PKM2 protein from human T cells stimulated with αCD3/αCD28 coated microbeads. Representative of two healthy donors. e) *PKM* isoform expression in human T cells stimulated as in **d**. Restriction digest of cDNA using *NcoI* (N), *PstI* (P), or both (NP) allowed for determination of *PKM2* to *PKM1* ratios (M2/M1) by dividing band intensity in the P lane by N lane^27^. Values are mean M2/M1 ratio ± SD of 3 healthy donors. f) *Pkm2* to *Pkm1* ratio determined by qRT-PCR in ARS2^f/f^ or ARS2^KO^ T cells following activation with αCD3/αCD28 + rIL-2 for indicated times (left) and total *Pkm* expression 3 days post activation (right). Graphs show mean ± SD, dots represent individual mice. Unpaired t test, n.s.= not significant, **p < 0.01.

We reasoned that splicing events in transcripts that were upregulated by T cell activation would be the most likely to have a biological consequence. We therefore examined transcripts that underwent ARS2-depdendent alternative splicing and were upregulated by T cell activation (Fig. 2c). Several interesting transcripts emerged from this analysis including *Pkm*, the transcript coding pyruvate kinase (PKM), the final enzyme of glycolysis. PKM is expressed as two isoforms that result from mutually exclusive alternative splicing; PKM1 is mainly expressed in quiescent cells, while PKM2 has been linked to the Warburg effect in tumor cells^25^. Consistent with mutually exclusive splicing of an upregulated transcript, human T cell activation induced PKM2 protein expression while having a limited effect on PKM1 protein (Fig. 2d). Similarly, PKM2 protein was enriched 3 days following *in vitro* or *in vivo* antigen stimulation of OT-I T cells (Extended Data Fig. 2a^26^). An established RT-PCR/restriction digest method^27, 28^ confirmed enrichment of *PKM2* mRNA relative to *PKM1* mRNA in activated human T cells (Fig. 2e). Isoform-specific qRT-PCR confirmed RNA-seq findings and showed that activated T cells preferentially induced *Pkm2* expression on day 3 in an ARS2-dependent manner (Fig. 2f, left). Importantly however, day 3 activated control and ARS2^KO^ T cells expressed equivalent total levels of *Pkm* mRNA (Fig. 2f, right), further indicating that ARS2 facilitates T cell activation induced *Pkm2* splicing.

### ARS2 recruits splicing factors to *Pkm* pre-mRNA

The mutually exclusive splicing of *Pkm* exons 9 and 10 is regulated by RNA binding proteins SRSF3, hnRNPA1/A2, and PTBP1^8^. Published RNA interactome data^29, 30, 31^ indicated ARS2, SRSF3, hnRNPA1, and PTBP1 all bound the *PKM* transcript in human cell lines, with ARS2, SRSF3, and PTBP1 peaks overlapping in exon 10 (Fig. 3a). RNA-immunoprecipitation (RIP) followed by qRT-PCR using intron-exon junction spanning primer sets to detect *Pkm* pre-mRNA confirmed these interactions in the murine hematopoietic cell line FL5.12 (Fig. 3b). Notably, 60 – 80% of *Pkm* pre-mRNA was pulled down with ARS2 and this interaction was disrupted by CBP80 knockdown (Fig. 3c and Extended Data Fig. 2b), indicating that ARS2 association with *Pkm* pre-mRNA was mediated by the CBC. Since the primary function of ARS2 bound to the CBC is recruitment of RNA processing factors to nascent transcripts^11, 12, 17, 24, 29^, we determined the effect of ARS2 on splicing factor association with *Pkm* pre-mRNA. For these experiments T cell activation was modeled by IL-3 stimulation of FL5.12.xL cells as previously described^32^. Similar to T cell activation, IL-3 stimulated FL5.12.xL cells upregulated ARS2 and preferentially spliced *Pkm* to favor *Pkm2* expression (Extended Data Fig. 2c,d). ARS2 knockdown during IL-3 stimulation of FL5.12.xL cells reduced binding of SRSF3 and hnRNPA1 to *Pkm* pre-mRNA (Fig. 3d and Extended Data Fig. 2e). These data inform a model wherein ARS2 upregulation by CD28 stimulates recruitment of splicing factors to *Pkm* pre-mRNA, thereby favoring splicing to *Pkm2* in activated T cells.

**Figure 3:**
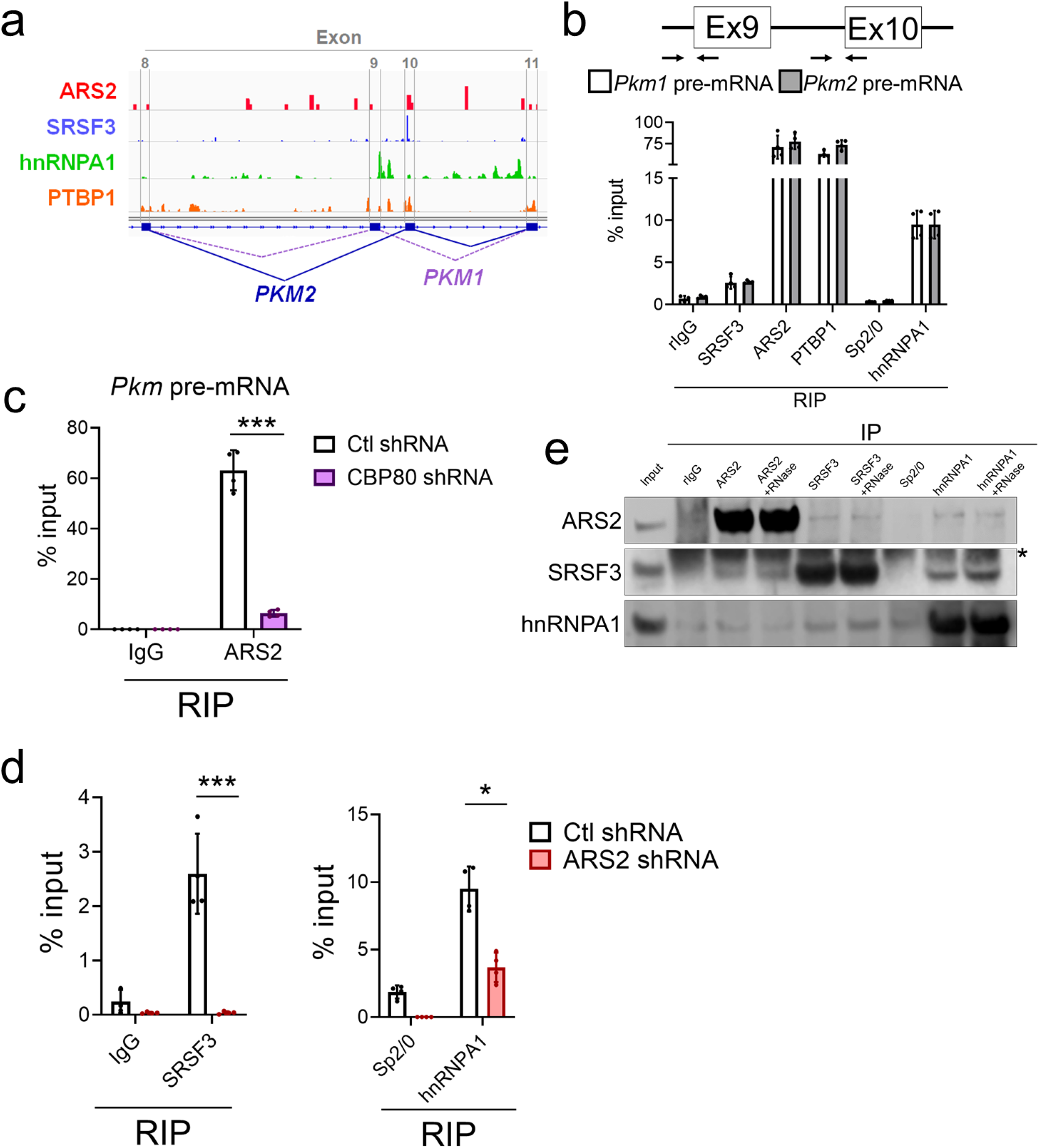
Recruitment of splicing factors to *Pkm* pre-mRNA by ARS2. a) RNA binding sites identified by crosslinking immunoprecipitation and sequencing for ARS2 (GSE94427), SRSF3 (GSE118265), hnRNPA1 (ENCSR154HRN), and PTBP1 (ENCSR981WKN) in HEK 293T cells mapped to exons 8 through 11 of the human *PKM* transcript. b) RNA immunoprecipitation (RIP) from FL5.12 cells using antibodies to indicated proteins or control antibodies (rIgG & Sp2/0) followed by qRT-PCR to detect *Pkm1* or *Pkm2* pre-mRNA with PCR primers spanning intron-exon boundaries as depicted. Graph shows mean ± SD of technical replicates from a representative experiment repeated twice. c) ARS2 RIP from CBP80 knockdown vs. control shRNA expressing FL5.12 cells. Bound *Pkm* pre-mRNA was quantified by qRT-PCR as in **b**. Graph shows mean ± SD of technical replicates from arepresentative experiment repeated twice. Unpaired t-test, n.s.= not significant, ***p < 0.001. d) SRSF3 and hnRNPA1 RIPs from ARS2 knockdown vs. control shRNA expressing FL5.12.xL cells stimulated with IL-3 for three days. Bound *Pkm* pre-mRNA was quantified as in **b**. Graphs show mean ± SD of technical replicates from representative experiment repeated twice. Unpaired t-test, n.s.= not significant, *p < 0.05, ***p < 0.001, ****p<0.0001. e) Co-immunoprecipitation (co-IP) identified protein-protein interactions between ARS2, SRSF3, and hnRNPA1. RNAse A was added to indicated lysates prior to IP. Input = 1% of total lysate. Replicate co-IP experiments are shown in Extended Data Fig. 2g. *light chain of IP antibody.

ARS2 interactome studies^12, 13, 24^ identified several splicing factors immunoprecipitated with ARS2 in human cell lines including SRSF3, hnRNPA1 and PTBP1 (Extended Data Fig. 2f). Reciprocal co-immunoprecipitation (co-IP) experiments performed using IL-3 stimulated FL5.12.xL cells confirmed ARS2 interaction with SRSF3, and hnRNPA1 (Fig. 3e and Extended Data Fig. 2g). Similar co-IPs were unable to detect interaction of ARS2 and PTBP1 (data not shown). Interestingly, co-IPs showed that SRSF3 and hnRNPA1 interacted with each other as well as ARS2 raising the possibility that these proteins form a ternary complex that directs alternative splicing of *Pkm* pre-mRNA, and potentially many more transcripts, in activated immune cells.

### ARS2 supports glycolytic reprogramming of activated T cells

Limited *Pkm2* splicing in activated T cells lacking ARS2 was predicted to alter glycolysis. To test this prediction, Seahorse flux analysis was used to measure glycolysis (defined by glucose-induced change in extracellular acidification rate (ECAR)) and oxidative phosphorylation (OXPHOS: defined as oligomycin-induced change in oxygen consumption rate (OCR)) of ARS2^f/f^ and ARS2^KO^ T cells 1 or 3 days following activation (Fig. 4a and Extended Data Fig. 2h). OXPHOS was equivalent between control and ARS2^KO^ T cells at both 1- and 3-days following activation. Additionally, ARS2 deletion did not alter glycolysis over the first 24 hour of activation. In line with temporal ARS2 regulation of *Pkm2* splicing following T cell activation (Fig. 2d-f), increased glycolysis observed in activated T cells between days 1 and 3 was completely abrogated by ARS2 deletion (Fig. 4b).

**Figure 4:**
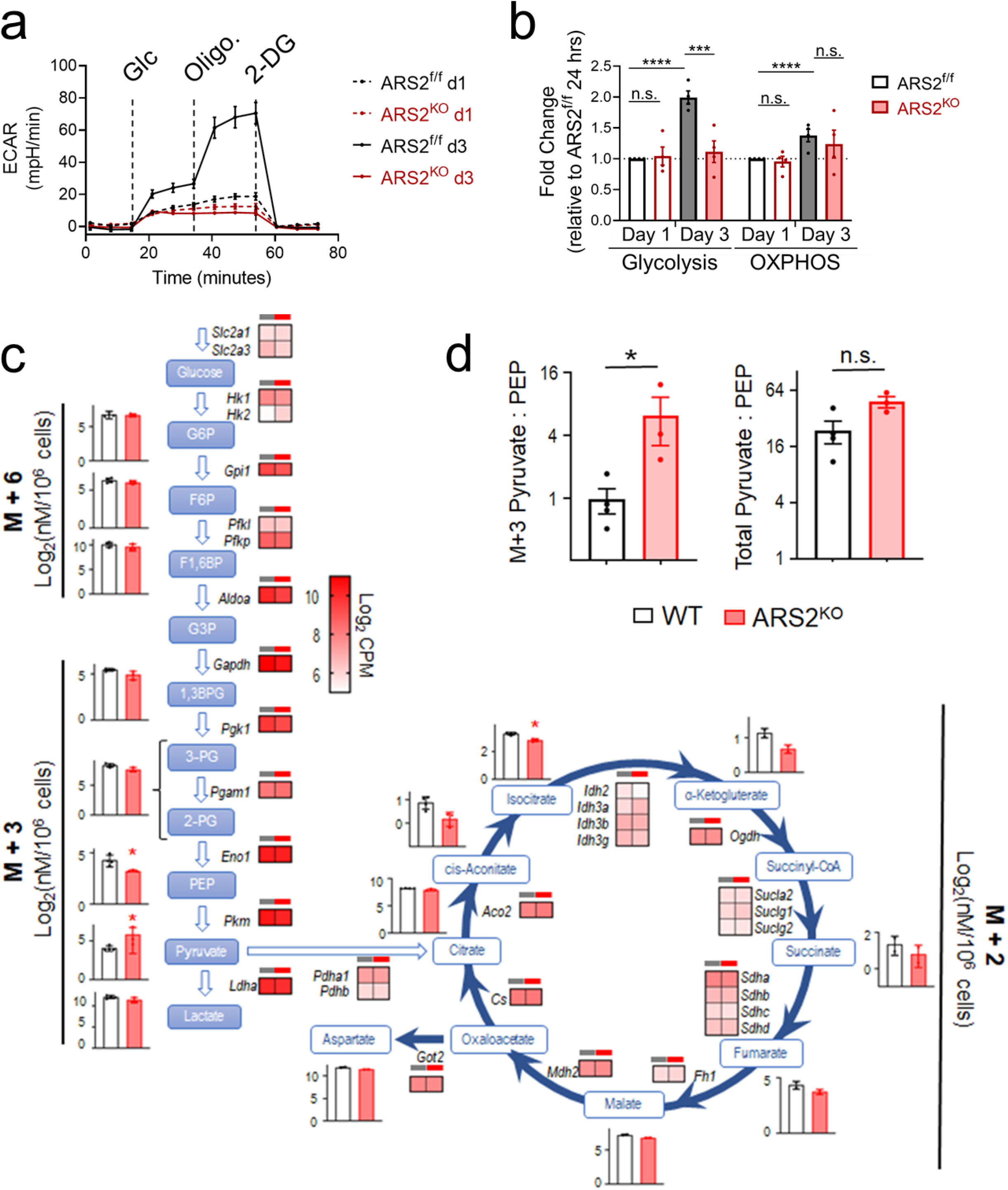
ARS2 supports T cell glycolytic reprogramming. a) Representative Seahorse Glycolysis Stress Test of ARS2^f/f^ and ARS2^KO^ T cells at 1 (dashed line) and 3 (solid line) days following activation with αCD3/αCD28 + rIL-2. Vertical lines indicate addition of glucose (Glc), oligomycin (Oligo.) and 2-deoxyglucose (2-DG). b) Quantification of glycolysis, defined as the change in ECAR following glucose (Glc) addition, and OXPHOS, defined as change in OCR following oligomycin (oligo) addition (see Extended Data Fig. 2h). Bars represent mean ± SD of 4 biological replicates, dots represent individual mice. Unpaired t-tests, n.s.= not significant, **p < 0.01, ***p < 0.001, ****p < 0.0001. c) [U-^13^C] glucose enrichment in glycolytic intermediates and citric acid cycle (TCA) metabolites in WT (black) or ARS2^KO^ (red) T cells activated for 3 days with αCD3/αCD28 + rIL-2. Bar graphs show mean ± SD of indicated isotopomers of each metabolite measured in 3 biological replicates. Unpaired t tests, *p < 0.05. Heatmaps show expression of indicated genes coding metabolic enzymes. d) Ratio of M+3 pyruvate to M+3 PEP and total pyruvate to total PEP in samples from **c**. Unpaired t test, n.s.= not significant, *p < 0.05.

To further characterize glycolysis in activated ARS2^KO^ CD8 T cells [U-^13^C] glucose tracing was employed. Tracing data and RNA-seq heatmaps obtained from WT and ARS2^KO^ CD8 T cells activated for 3 days were overlaid onto a diagram of central carbon metabolism to visualize changes resulting from loss of ARS2 (Fig. 4c). Consistent with the small decrease in glucose uptake observed (Extended Data Fig. 2i), expression of GLUT3 (*Slc2a3*) was reduced in ARS2^KO^ T cells. Increased expression of hexokinase (*Hk2*) may compensate for reduced GLUT3 by allowing ARS2^KO^ T cells to phosphorylate glucose more efficiently, leading to equivalent ^13^C-labeling of glucose 6-phosphate (^13^C_6_ or M+6 G6P). All other transcripts coding glycolysis enzymes were similar between WT and ARS2^KO^ T cells. ^13^C labeling of glycolytic intermediates was comparable until the final rate-limiting step of glycolysis, conversion of phosphoenolpyruvate (PEP) to pyruvate, where we observed increased pyruvate labeling (^13^C_3_ or M+3) and decreased PEP labeling (^13^C_3_ or M+3) leading to an increased ratio of labeled (^13^C_3_ or M+3) pyruvate to PEP in ARS2^KO^ T cells (Fig. 4d). The ratio of total (^13^C-labeled + unlabeled) pyruvate to total PEP was much higher than the ratio of M+3 pyruvate to M+3 PEP and was only slightly altered by loss of ARS2, suggesting that a non-glycolytic source of pyruvate exists in activated T cells. Overall, metabolic data show that ARS2 influences pyruvate kinase activity in support of continued glycolytic reprogramming following the well-characterized induction of glycolysis during T cell priming.

### PKM2 supports CD8 T cell anti-tumor immunity

To directly test the function of PKM2 in activated CD8 T cells, we generated *Pkm2* inducible knockout mice (PKM2^iKO^) by crossing *Pkm2*^f/f^ mice^33^ to CreERT2 expressing mice^34^. Tamoxifen administration to PKM2^iKO^ mice led to loss of PKM2 from peripheral T cells and compensatory upregulation of PKM1 (Extended Data Fig. 3a,b). Loss of PKM2 did not alter the number or frequency of splenic T cells (Extended Data Fig. 3c) nor did it reduce their ability to express activation markers following stimulation with αCD3/αCD28 + rIL-2 (Extended Data Fig. 3d). Seahorse assays demonstrated that glycolytic reprogramming of activated PKM2^KO^ T cells was similar to ARS2^KO^ T cells, with a lack of increased glycolysis between day 1 and day 3 seen in both (Fig. 5a,b and Extended Data Fig. 3e). Functionally, adoptively transferred OT-I transduced PKM2^KO^ CD8 T cells were unable to control growth of E.G7-OVA tumors as well as WT controls (Fig. 5c and Extended Data Fig. 3f). Furthermore, PKM2^KO^ effector CD8 T cells were unable to fully induce IFNγ protein expression despite exhibiting similar expression of *Ifng* mRNA (Fig. 4d,e). Observed defects in PKM2^KO^ T cells closely mirror those observed in ARS2^KO^ T cells and support the notion that ARS2-directed alternative splicing to favor PKM2 expression is a mechanism by which ARS2 influences anti-tumor T cell immunity.

**Figure 5:**
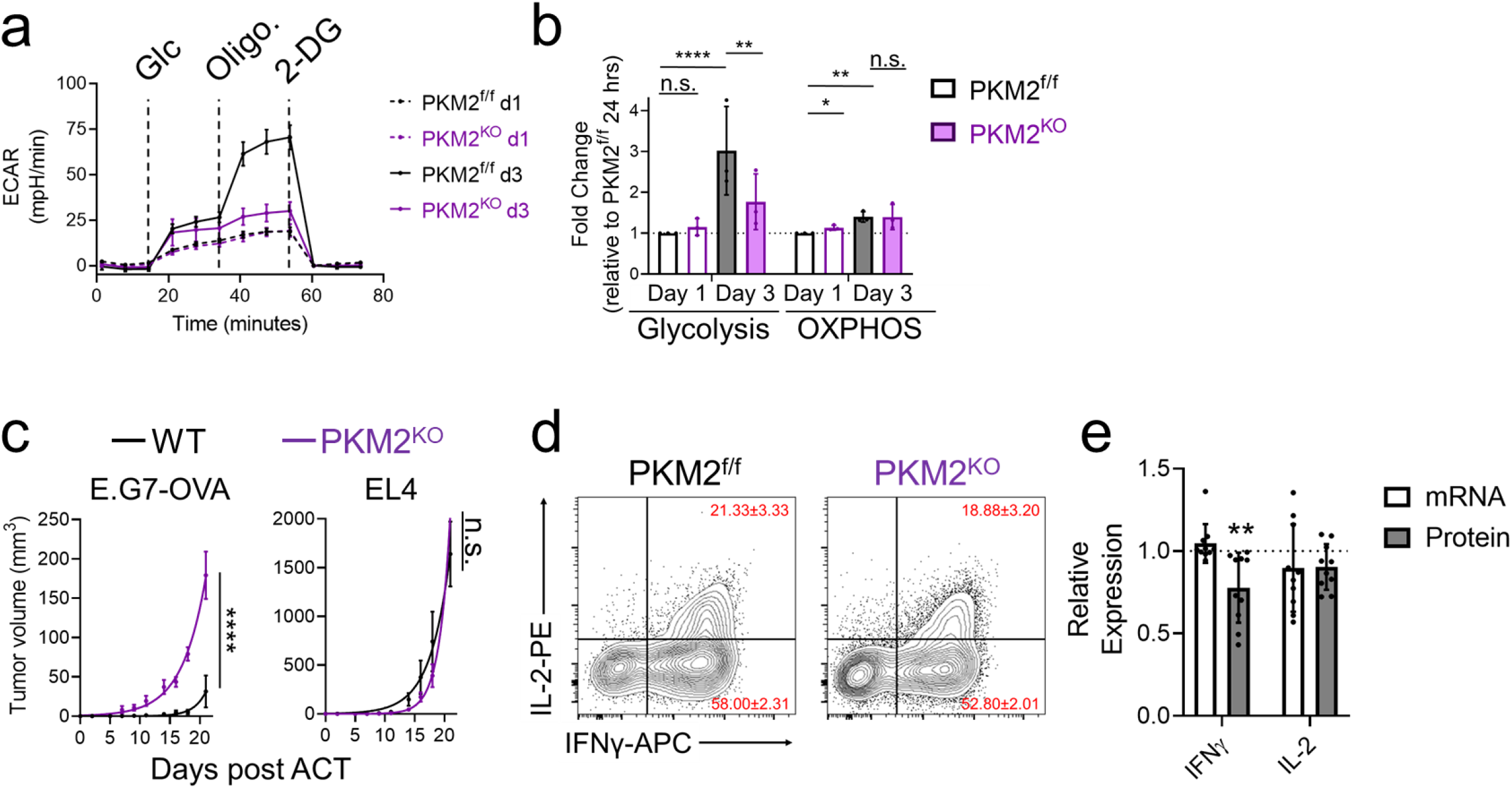
PKM2 supports glycolytic reprogramming and CD8 T cell effector function. a) Representative Seahorse Glycolysis Stress Test of PKM2^f/f^ and PKM2^KO^ T cells at 1 (dashed line) and 3 (solid line) days post-activation. Vertical lines indicate addition of glucose (Glc), oligomycin (Oligo.) and 2-deoxyglucose (2-DG). b) Quantification of glycolysis and OXPHOS (see Extended Data Fig. 3e) of activated PKM2^f/f^ and PKM2^KO^ T cells. Bars represent mean ± SD of 4 biological replicates, dots represent individual mice. Unpaired t-tests, n.s.= not significant, *p < 0.05, **p < 0.01, ****p < 0.0001. c) Growth of E.G7-OVA and EL4 tumors on opposite flanks of C57BL/6J mice following adoptive transfer of CD8 T cells engineered to express the OT-I TCR. Graphs show best fit lines to mean tumor volume ± SEM in mice receiving OT-I transduced WT (n=8) or PKM2^KO^ (n=6) T cells. Two-way ANOVA, n.s.= not significant, ****p < 0.0001. d) Flow cytometry assessment of IFNγ and IL-2 expression in activated PKM2^fl/fl^ or PKM2^KO^ effector T cells (CD8^+^CD44^+^CD62L^-^) following re-stimulation with PMA/ionomycin in the presence of brefeldin-A. Red text indicates mean percent of cells found in each region ± SD of 4 biological replicates. e) IFNγ and IL-2 protein (from **d**) and mRNA expression in PKM2^KO^ T cells relative to control PKM2^fl/fl^ T cells. Graphs show mean ± SD; dots represent individual mice. One-way ANOVA, **p < 0.01, ***p < 0.001.

### CD28 PYAP signaling drives ARS2-dependent *Pkm2* splicing

Having established that ARS2-dependent *Pkm2* alternative splicing drives glycolytic metabolism and anti-tumor function of activated CD8 T cells, we sought to further elucidate the T cell activation signals regulating this pathway. CD28 activates signaling through at least three intracellular domains (ICDs) in its cytoplasmic tail^35^, the membrane proximal YMNM ICD that activates phosphoinositide 3-kinase (PI3K) signaling, the membrane distal PYAP ICD that activates signaling through lymphocyte-specific protein tyrosine kinase (Lck) and growth factor receptor-bound protein 2 (Grb2), and the PRRP ICD that lies between the YMNM and PYAP ICDs and activates signaling through interleukin-2-inducible kinase (Itk; Fig. 6a). To examine contributions of each domain to T cell activation-induced ARS2 upregulation, we stimulated WT, CD28 knockout^36^ (CD28^KO^), CD28^AYAA^ knock-in^37^, CD28^Y170F^ knock-in^38^, and CD28 AYAA and Y170F double knock-in^39^ (CD28^DKI^) T cells with αCD3/αCD28 + rIL-2 to compensate for decreased IL-2 production by CD28 mutant T cells^37^. These experiments revealed that ARS2 upregulation required CD28 signaling through its PYAP ICD, while signaling through the CD28 PRRP ICD was required to maintain basal levels of ARS2 expression (Fig. 6b). Importantly, CD28 mutant T cells expressed similar basal levels of ARS2 and were capable of ARS2 upregulation following stimulation with PMA and ionomycin (Extended Data Fig. 4a,b).

**Figure 6:**
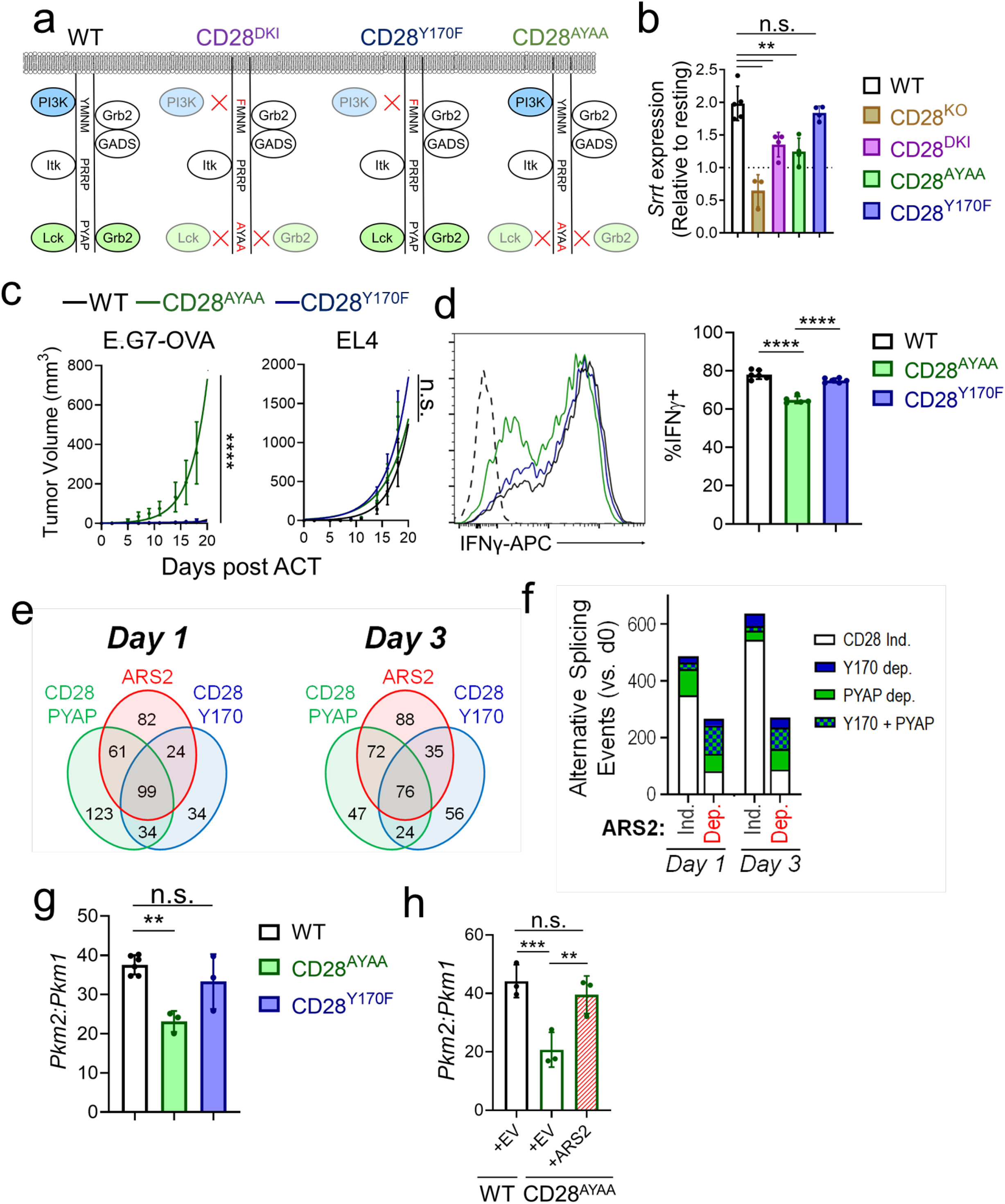
CD28 signaling drives ARS2-dependent alternative splicing. a) Diagram of CD28 intracellular domains and associated signaling proteins. Knock-in mice express CD28 with intracellular domains containing loss of function mutations indicated in red. b) ARS2 mRNA (*Srrt*) expression in WT and CD28 mutant T cells stimulated with αCD3/αCD28 + rIL-2 for 24 hours. Graphs show mean ± SD. Dots represent individual mice. One-way ANOVA, n.s.= not significant, **p < 0.01. c) Growth of E.G7-OVA and EL4 tumors on opposite flanks of C57BL/6J mice following adoptive transfer of CD8 T cells engineered to express the OT-I TCR. Graphs show best fit lines to mean tumor volume ± SEM in mice receiving OT-I transduced WT (n = 8), CD28^AYAA^ (n = 5), or CD28^Y170F^ (n = 5) T cells. Two-way ANOVA, n.s.= not significant, ****p < 0.0001. d) Flow cytometry assessment of IFNγ expression in CD28 mutant effector T cells (CD8^+^CD44^+^CD62L^-^) following re-stimulation with PMA/ionomycin in the presence of brefeldin-A. Representative histograms are shown (dashed line = FMO control). Bar graphs show mean ± SD of biological replicates, dots represent individual mice. One-way ANOVA, ****p<0.001. e) Venn diagrams showing overlap of AS events dependent on ARS2 (red), CD28 -PYAP (green) and CD28-PI3K (blue) at day 1 (left) and day 3 (right) post-activation. f) Quantification of AS events that are ARS2 dependent and also show dependency on CD28 signaling through either its PYAP ICD, YMNM ICD, or both at day 1 and day 3 post -activation. g) Ratio of *Pkm2* to *Pkm1* in WT and CD28 mutant T cells 3 days post activation with αCD3/αCD28 + rIL-2. Graphs show mean ± SD, dots represent individual mice. One-way ANOVA, **p<0.01, ***p<0.001. h) WT or CD28^AYAA^ T cells were isolated, activated with αCD3/αCD28 + rIL-2, and transduced with an empty retroviral vector (EV) or a retroviral ARS2 expression vector. Ratio of *Pkm2* to *Pkm1* measured by qRT-PCR on day 3 post activation is shown. Graphs show mean ± SD, dots represent individual mice. One-way ANOVA, *p < 0.05, **p < 0.01.

The CD28 PYAP ICD necessary for ARS2 upregulation was previously shown important for *in vivo* T cell function in models of allergic airway inflammation and experimental autoimmune encephalomyelitis ^37, 38, 39^, both of which are primarily driven by CD4 T cells. To test whether the CD28 PYAP domain was also important for CD8 T cell anti-tumor immunity, OT-I TCR transduced WT, CD28^AYAA^ knock-in and CD28^Y170F^ knock-in CD8 T cells were adoptively transferred into E.G7-OVA/EL4 tumor bearing mice as in Fig. 1g. Similar to what was seen using ARS2^KO^ and PKM2^KO^ OT-I transduced CD8 T cells, adoptive transfer of OT-I transduced CD28^AYAA^ CD8 T cells did not control E.G7-OVA tumor growth as well as WT controls, while OT-I transduced CD28^Y170F^ CD8 T cells demonstrated anti-tumor function similar to WT controls (Fig. 6c and Extended Data Fig. 4c,d). IFNγ expression in CD28^AYAA^ CD8 effector T cells was also reduced compared to WT or CD28^Y710F^ knock-ins (Fig. 6d and Extended Data Fig. 4d).

Since functional data were consistent with a CD28-ARS2 pathway activated by CD28 PYAP ICD signaling we repeated RNA-seq analysis of T cell activation-induced alternative splicing using CD28 mutant T cells. PSI-sigma analysis revealed marked overlap between T cell activation-induced alternative splicing events regulated by CD28 ICDs and ARS2 (Fig. 6e), with more than half of ARS2 dependent splicing events also requiring intact CD28 PYAP signaling. In contrast, the great majority of ARS2 independent splicing events occurred normally in the absence of CD28 signaling (Fig. 6f). Notably, RNA-seq demonstrated that *Pkm2* splicing was regulated by both ARS2 and CD28 PYAP signaling. Regulation of *Pkm2* splicing by CD28 PYAP signaling was confirmed by qRT-PCR (Fig. 6g). Importantly, the reduced ratio of *Pkm2* to *Pkm1* resulting from defective alternative splicing in CD28^AYAA^ T cells was rescued to WT level by retroviral transduction with ARS2 (Fig. 6h and Extended Data Fig. 4f), confirming that ARS2 upregulation by CD28 PYAP signaling induced *Pkm2* splicing.

### The CD28-ARS2-PKM2 axis licenses T cell glycolysis and IFNγ production

Since loss of ARS2-dependent *Pkm2* splicing resulted in defective glycolysis 3 days following T cell activation, we repeated Seahorse assays and [U-^13^C] glucose tracing using CD28 knock-in T cells to determine if upstream CD28 signaling resulted in similar downstream glycolytic phenotypes to those observed in ARS2^KO^ and/or PKM2^KO^ T cells. Activated CD28^AYAA^ T cells phenocopied ARS2^KO^ and PKM2^KO^ T cells is Seahorse assays, demonstrating reduced glycolysis and no changes in OXPHOS relative to WT controls (Fig. 7a,b and Extended Data Fig. 4g). Additionally, the ratio M+3 labeled pyruvate to M+3 PEP in day 3 activated CD28^AYAA^ T cells looked very similar to ARS2^KO^ T cells, although differences compared to WT did not reach statistical significance (Fig. 7c and Extended Data 4h).

**Figure 7:**
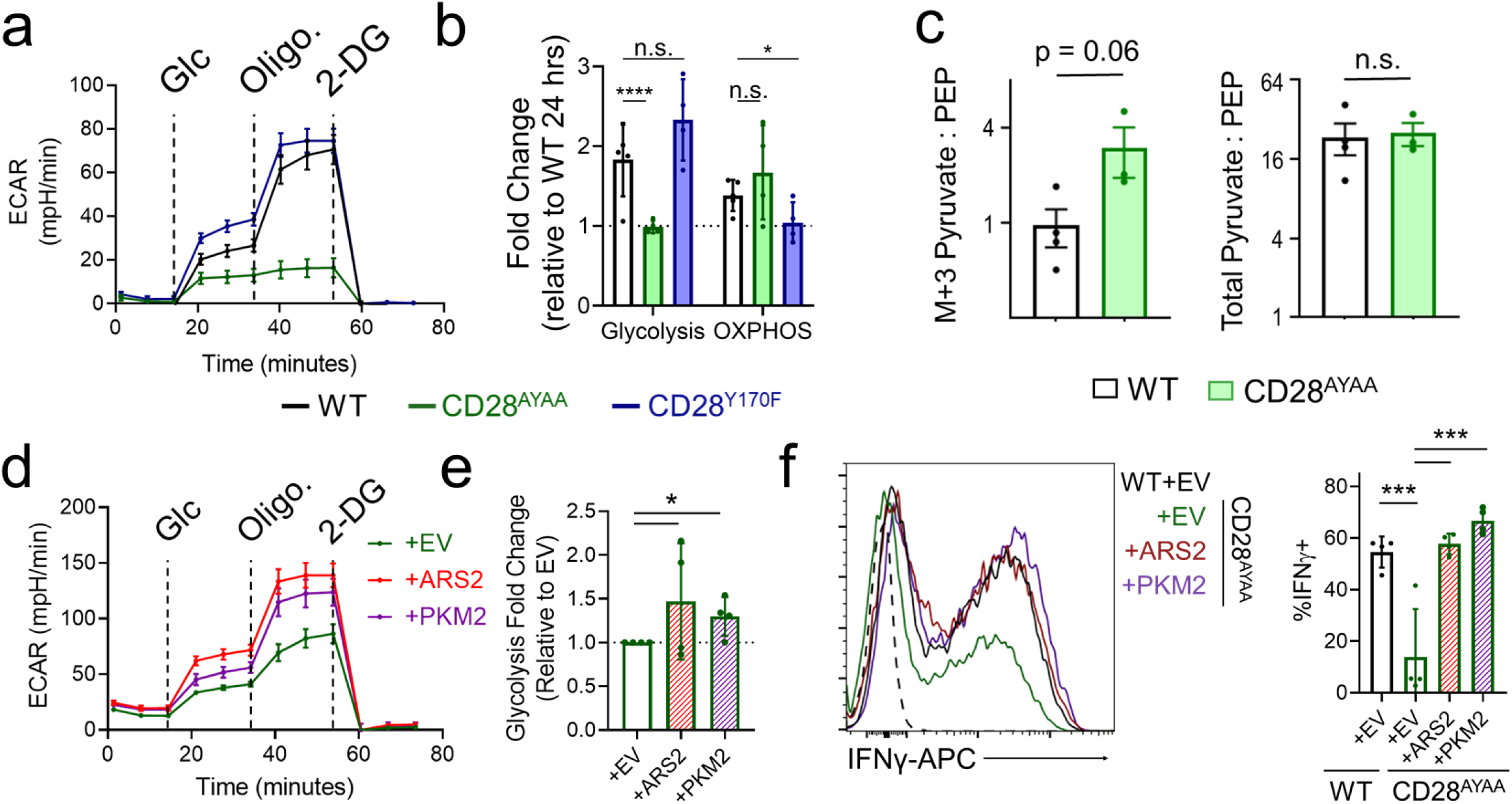
The CD28-ARS2-PKM2 axis supports glycolytic metabolism and effector cytokine production. a) Representative Seahorse Glycolysis Stress Test of WT and CD28 mutant T cells 3 days following stimulation with αCD3/αCD28 + rIL-2. Vertical lines indicate addition of glucose (Glc), oligomycin (Oligo.) and 2-deoxyglucose (2-DG). b) Quantification of glycolysis and OXPHOS (see Extended Data Fig. 4g) in T cells described in **a**. Bars represent mean ± SD of 4 biological replicates, dots represent individual mice. One-way ANOVA, n.s.= not significant, **p < 0.01, ***p < 0.001. c) Ratio of M+3 pyruvate to M+3 PEP and total pyruvate to PEP in cells described in **a** incubated with [U-^13^C] glucose. Bar graphs show mean ± SD of biological replicates, dots represent individual mice, unpaired t test, n.s.= not significant. d) CD28^AYAA^ T cells were isolated, activated with αCD3/αCD28 + rIL-2, and transduced with an empty retroviral (EV) or retroviral ARS2 or PKM2 expression vectors. Representative Seahorse Glycolysis Stress Test preformed on transduced T cells 3 days following activation is shown. Vertical lines indicate addition of glucose (Glc), oligomycin (Oligo.) and 2-deoxyglucose (2-DG). e) Quantification of glycolysis in T cells described in **d**. Bars represent mean ± SD of 4 biological replicates, dots represent individual mice. One-way ANOVA, *p < 0.05. f)Flow cytometry assessment of IFNγ expression in effector T cells (CD8 ^+^CD44^+^CD62L^-^) described in **d** following re-stimulation with PMA/ionomycin in the presence of brefeldin-A. Representative histograms are shown (dashed line = FMO control). Bar graphs show mean ± SD of biological replicates, dots represent individual mice. Unpaired t test, ***p < 0.001.

Rescue experiments were performed to directly test whether ARS2-dependent *Pkm2* alternative splicing contributed to defective T cell glycolysis observed in activated CD28^AYAA^ T cells. Retroviral transduction of CD28^AYAA^ T cells with ARS2 or PKM2 expression constructs led to increased glycolysis 3 days following activation when compared to empty vector (EV) controls (Fig. 7d,e). Notably, *Pkm2* to *Pkm1* ratio was similar to WT in ARS2 and PKM2 transduced CD28^AYAA^ T cells (Extended Data Fig. 4f). Since glycolysis is a known driver of *Ifng* mRNA translation^4^, we further examined the ability of ARS2 and PKM2 to rescue IFNγ production in CD28^AYAA^ T cells. Retroviral transduction with an empty vector reduced the frequency of WT or CD28^AYAA^ CD8 effector T cells that express IFNγ in response to PMA + ionomycin re-stimulation (Fig. 7f vs. Fig. 6d). Despite this general reduction, rescue of CD28^AYAA^ T cells with ARS2 or PKM2 resulted in IFNγ expression similar to empty vector transduced WT T cells (Fig. 7f).

In summary, data presented here inform a mechanism whereby CD28 co-stimulation induces signaling through its membrane-distal PYAP intracellular domain that results in upregulation of ARS2, which in turn induces *Pkm2* splicing to enable CD8 T cell glycolytic metabolism, effector cytokines production, and anti-tumor immunity.

## Discussion

To provide protective anti-tumor immunity, naïve tumor antigen specific CD8 T cells must dramatically remodel their transcriptomes upon antigen encounter to drive cellular changes underlying effector differentiation and function. While the role of transcription factors in this process has been extensively characterized much less is known about how co-transcriptional processes, such as RNA splicing, contribute to acquisition of CD8 T cell effector properties. Studies defining roles for alternative splicing in T cells have largely focused on CD4 T cells, where splicing of surface receptors (*CD45, CD44*), signaling molecules (*MALT1*) and transcription factors (*FOXP3*) all contribute to CD4 differentiation and/or function^9^. Additionally, splicing factors including hnRNPLL and CELF2 were shown to be upregulated by TCR and/or CD28 co-receptor signaling in CD4 T cells and to influence T cell activation induced alternative splicing^40, 41^. A recent study found that alternative splicing patterns were similar between CD4 and CD8 T cells^42^, yet regulation and functional implications of alternative splicing in CD8 T cells remains mostly unexplored. The current study addresses this gap in knowledge by showing that the CD28 regulated expression of the CBC adaptor protein ARS2 regulates splicing of approximately one-third of T cell activation-induced alternative splicing events, including mutually exclusive splicing to favor expression of the M2 isoform of PKM. Furthermore, the novel CD28-ARS2-PKM2 axis stimulates continued reprogramming of glycolysis in activated CD8 T cells to support effector cytokine production and anti-tumor immunity.

ARS2 has emerged over the past decade as a critical intermediary physically linking the CBC to RNA maturation and degradation pathways long known to rely on RNA capping and CBC binding ^10^. Data presented here are, to the best of our knowledge, the first to demonstrate that ARS2 regulates the CBC-dependent process of alternative RNA splicing. Coupled with data demonstrating physical interactions between SRSF3, hnRNPA1, and ARS2 that promotes SRSF3 and hnRNPA1 association with *Pkm* pre-mRNA, we provide compelling mechanistic evidence that ARS2 facilitates alternative splicing during T cell activation. Data clearly indicate that ARS2 regulates many additional splicing events during T cell activation, potentially by recruiting SRSF3, hnRNPA1, and/or other splicing factors to nascent transcripts. Further examination of ARS2-directed alternative splicing events, or potentially other ARS2-directed RNA maturation events, may reveal additional ways in which ARS2 contributes to T cell biology. Moreover, it is unclear if the newly identified splicing function for ARS2 is limited to activated immune cells or is more broadly relevant, as is if regulation of alternative splicing by ARS2 underlies its essential role in thymic T cell development^15^.

Knockout experiments established that one product of ARS2-directed alternative splicing, PKM2, was required for activated CD8 T cells to fully induce glycolysis, produce high levels of IFNγ, and mediate efficient antigen specific control of tumor growth in mice. Glycolysis and IFNγ expression were similarly reduced in CD4 T cells by PKM2 inhibition^43^ or knockout^44^, yet in CD4 T cells PKM2 appeared to have a more dominant role in directing Th17 differentiation independent of its metabolic function^5, 6, 7^. However, the non-metabolic role of PKM2 in directly driving transcription remains controversial^8^ and experiments to tease out metabolic versus non-metabolic functions of PKM2 are quite challenging, especially in light of the growing body of evidence that metabolic perturbations often directly influence the function of epigenetic enzymes^45^. Our data indicate PKM2 drives IFNγ production in CD8 effector T cells, and perhaps IFNγ-dependent anti-tumor immunity, post-transcriptionally and therefore most likely through induction of glycolysis and resulting relief of GAPDH repression of *Ifng* mRNA translation^4^. Whether non-metabolic functions of PKM2 contribute to CD8 T cell anti-tumor activity remains to be determined.

We recently reported that signaling through the CD28 PYAP ICD is required for enhanced glycolysis in T cells activated for 24 hours^46^. Earlier studies found that PI3K activation by CD28 induced GLUT1 expression and enhanced glycolysis over the same 24 hour activation period^3, 47^, indicating that the signaling initiated by the membrane proximal YMNM and membrane distal PYAP domains of CD28 cooperate to induce glycolytic metabolism during T cell priming. Here we find that only CD28 PYAP signaling is necessary for continued induction of glycolysis beyond levels observed during T cell priming, adding additional complexity to what is currently known about CD28 regulation of T cell metabolic reprogramming. Intuitively, it makes sense that metabolic states shift to meet cellular demands as T cells respond to activation stimuli with cell growth followed by proliferative expansion and differentiation into effector or memory populations. The mechanism elucidated by data presented here appears to almost entirely account for continued induction of glycolysis from days 1 to 3 of T cell activation and indicates that this phase of glycolytic reprogramming is necessary for optimal IFNγ production and anti-tumor immunity mediated by effector T cells. Developing strategies to target the CD28-ARS2-PKM2 axis identified here may therefore increase efficacy of cancer immunotherapy.

## Supporting information

Holling et al Supplemental Figures 1-4

## Acknowledgments

The authors thank members of the Lee lab for insightful discussion about the manuscript, Louise Carlson for maintaining CD28 mutant mice, Dr. Fumito Ito for providing PMEL mice for breeding, Jessie Chiello for running Seahorse assays, and Dr. Prashant Singh for consultation on RNA-seq experiments. This work was supported by National Cancer Institute grants R00CA175189 (to S.H.O.), R01CA205246 (to E.A.R.), R01CA121044 (to K.P.L.), T32CA085183 (to G.A.H), and P30CA016056 involving the use of the Roswell Park Comprehensive Cancer Center Flow and Image Cytometry, Genomics, Laboratory Animal, and Immune Analysis Shared Resources, and by the Roswell Park Alliance Foundation. NMR experiments were carried out at the Center for Environmental and Systems Biochemistry Shared Resource Facility funded in part by the Markey Cancer Center P30CA177558.

## Author contributions

Conceptualization, G.A.H. and S.H.O.; Methodology, G.A.H., G.Q., S.H.O; Software, A.P.S and K.H.E; Formal Analysis, G.A.H., A.P.S., K.H.E, S.H.O.; Investigation, G.A.H., M.M.H., C.M.J., S.L.M., G.Q., S.H., K.L.S., T.R.E., T.G.; Resources, B.H.S., A.M.I., T.W.M.F, A.N.L., R.M.H., E.A.R., K.P.L., S.H.O.; Data Curation, A.P.S.; Writing – Original Draft, G.A.H., S.H.O.; Visualization, G.A.H., A.P.S., S.H.O.; Supervision and Project Administration, S.H.O, Funding Acquisition, G.A.H., E.A.R, K.P.L, S.H.O.

