## Supplementary material for "ARS2-directed alternative splicing mediates CD28 driven T cell glycolysis and effector function": Holling et al Supplemental Figures 1-4

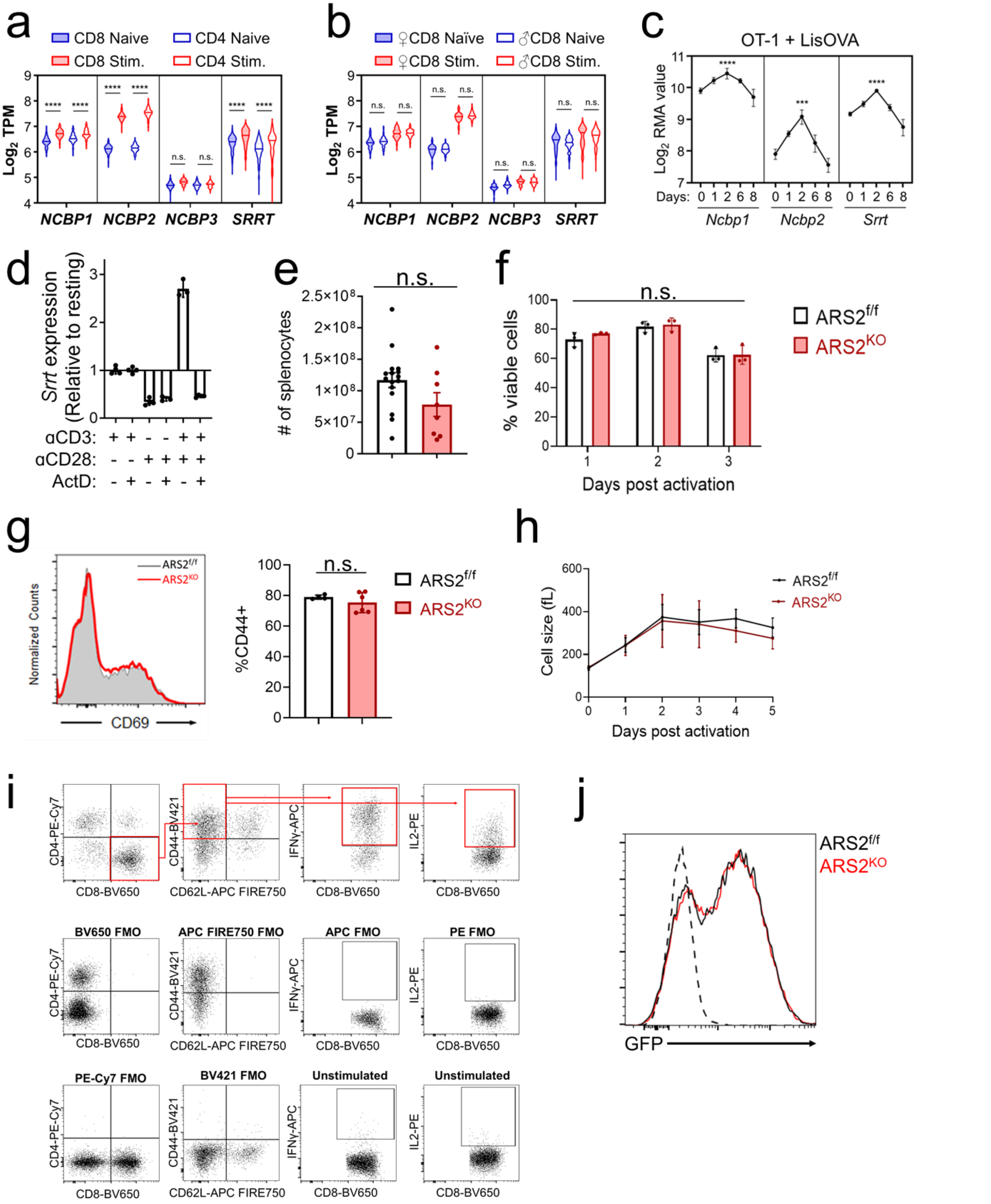


**Supplementary Figure 1:**

a) Violin plot of expression of CBCA components (and *NCBP3*) in CD4⁺ and CD8⁺ T cells (naïve and stimulated) from DICE database, (unpaired t test, n.s. = not significant, ****p<0.0001).

b) Expression of CBCA components stratified by male and female from DICE database, (unpaired t test, n.s. = not significant).
c) Expression of transcripts coding CBCA components in CD8 OT-I TCR transgenic T cells at indicated timepoints following infection of mice with LmOVA; from Immgen dataset GSE15907. Two-way ANOVA, ***p<0.001, ****p<0.0001.
d) ARS2 (*Srrt*) induction in WT T cells stimulated with either αCD3, αCD28, or both ± 1μg/mL Actinomycin D.
e) # of splenocytes found in ARS2$^{f/f}$ or ARS2$^{KO}$ mice following 5 days of tamoxifen treatment. Graphs show mean ± SD, dots represent individual mice (unpaired t test, n.s. = not significant).
f) Viability of ARS2$^{f/f}$ or ARS2$^{KO}$ T cells following stimulation with αCD3/αCD28 + rIL-2 measured by DAPI using flow cytometry. Graphs show mean ± SD, dots represent individual mice (unpaired t test, n.s. = not significant).
g) Representative flow plot of CD69 expression ARS2$^{f/f}$ or ARS2$^{KO}$ T cells 24 hours post-stimulation (left) and CD44 72 hours post-activation (right). Graph shows mean ± SD, dots represent individual mice (unpaired t test, n.s. = not significant).
h) Cell size of ARS2$^{f/f}$ or ARS2$^{KO}$ T cells following stimulation. Graphs show mean ± SD of 3 biological replicates.
i) Gating scheme and FMOs used to quantify IFNγ and IL-2 production of stimulated T cells.
j) GFP expression of OT-I transduced ARS2$^{f/f}$ or ARS2$^{KO}$ CD8$^{+}$ T cells (used in **Fig. 1g**).

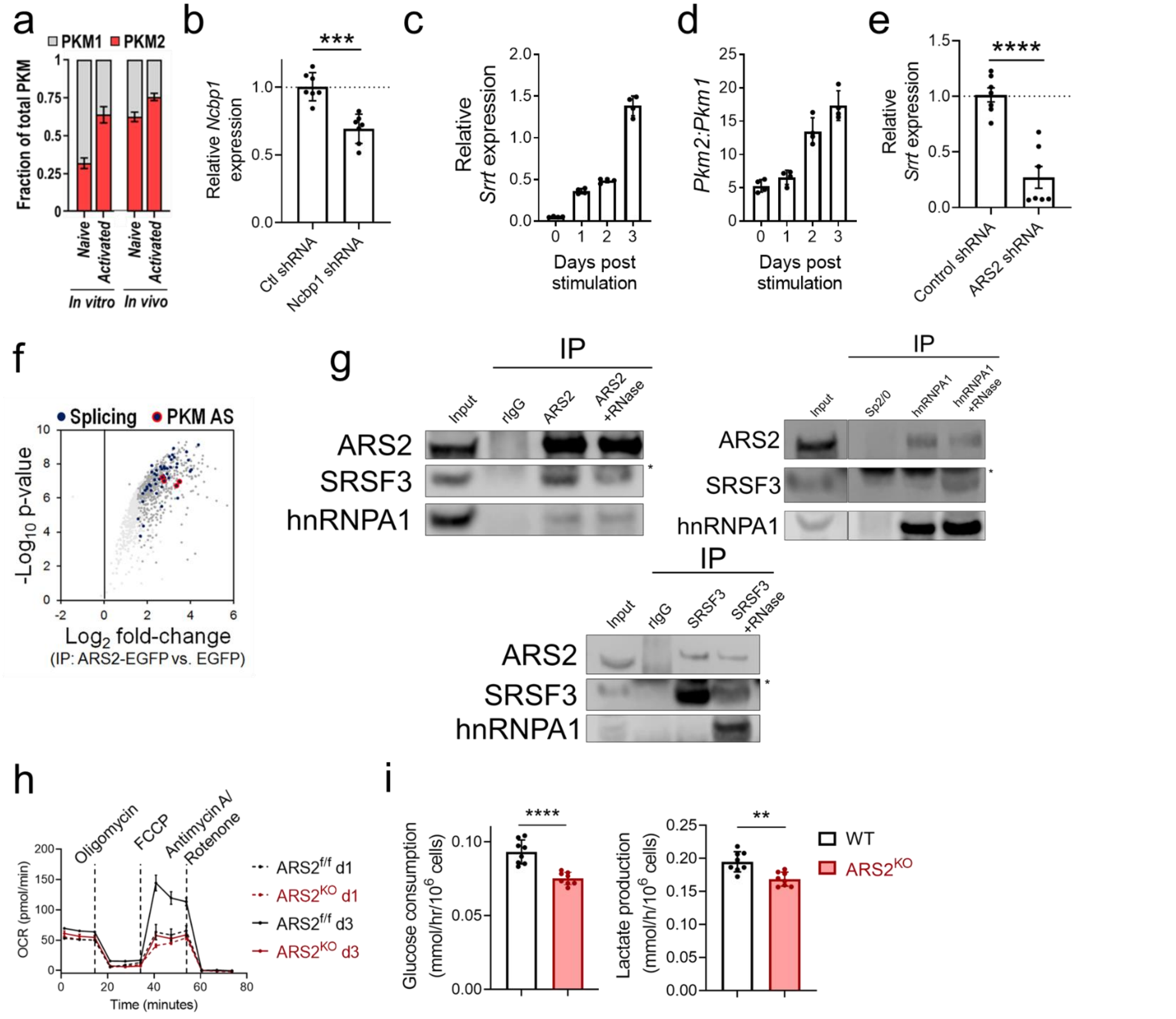


**Supplementary Figure 2:**

a) Fractional enrichment of PKM isoforms (publicly available data from Ma et al., 2019[26]).
b) Validation of CBP80 (*Ncbp1*) shRNA. Data shows mean ± SD of technical replicates from 2 independent experiments (unpaired t test, ***p<0.001).
c) ARS2 (*Srrt*) expression in FL5.12.xL subjected to IL-3 withdrawal for 5 days, followed by stimulation with IL-3 for 3 days.
d) Ratio of *Pkm2:Pkm1* in FL5.12.xL subjected to IL-3 withdrawal for 5 days, followed by stimulation with IL-3 for 3 days
e) Validation of ARS2 (*Srrt*) shRNA. Data shows mean ± SD of technical replicates from 2 independent experiments (unpaired t test, ****p<0.0001).
f) Interaction of ARS2 with known *Pkm* splicing factors from publicly available mass spectrometry data[12, 13, 24].
g) Confirmatory co-immunoprecipitations (related to Fig. **3e**). *light chain of IP antibody.
h) Representative Seahorse mitochondrial stress test of ARS2$^{f/f}$ or ARS2$^{KO}$ T cells at the indicated time point.
i) Glucose consumption (left) and lactate excretion (right) measured by T cells of indicated genotypes activated with αCD3/αCD28+rIL-2 three days earlier. Unpaired t test, **p<0.01, ****p<0.0001.

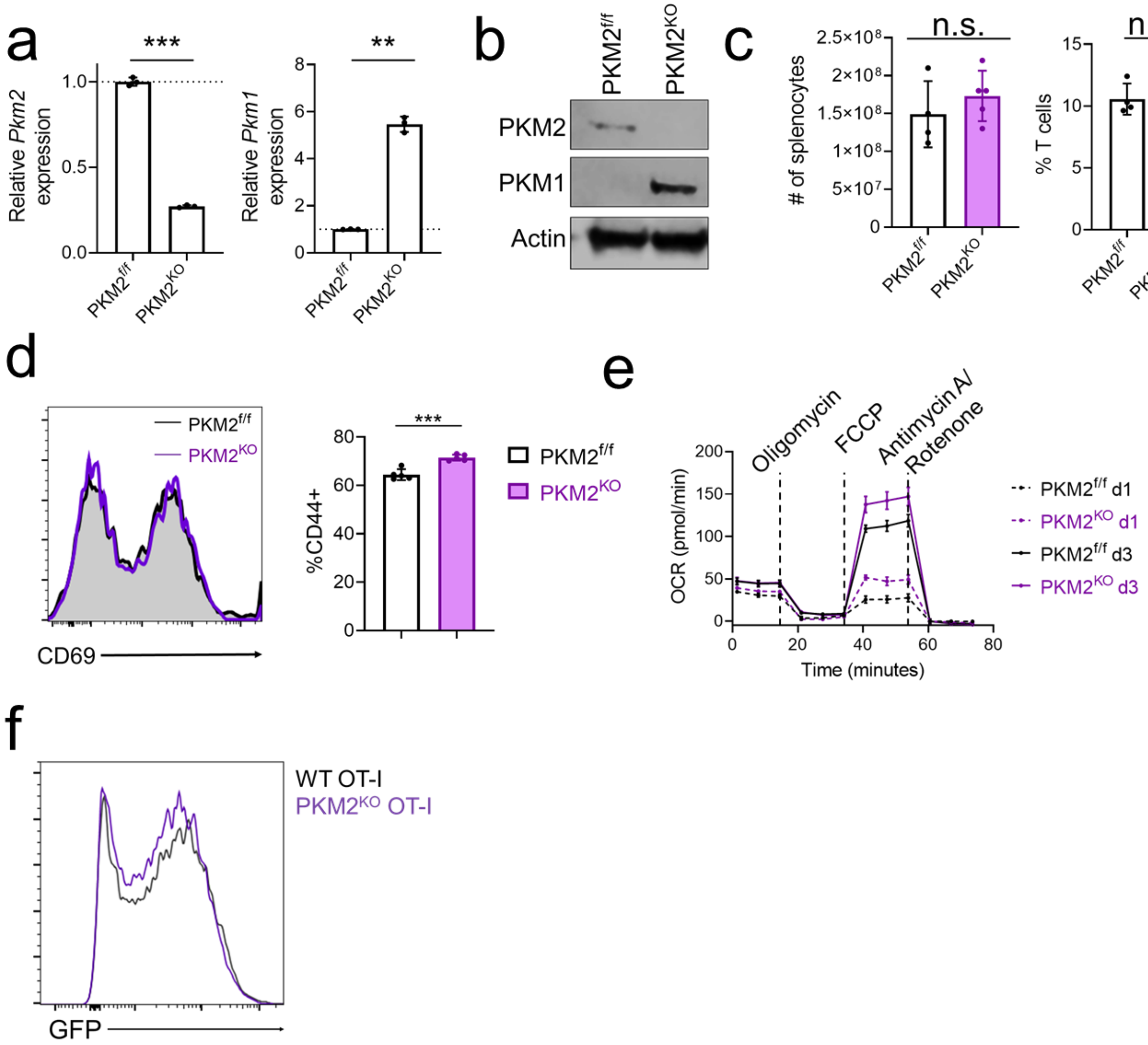


**Supplementary Figure 3:**

a) Validation of PKM2$^{iKO}$ model showing reduced expression of *Pkm2* (left) and increased expression of *Pkm1* (right) in purified T cells following 5 days of tamoxifen treatment. Graphs show mean ± SD, (unpaired t test, **p<0.01, ***p<0.001).
b) Same as in **a** but showing PKM1 and PKM2 protein expression.
c) # of splenocytes (left) and frequency of peripheral T cells (right) from PKM2$^{iKO}$ mice. Graphs show mean ± SD, each dot represents one mouse (unpaired t test, n.s. = not significant).
d) Representative flow plot of CD69 expression 24 hours post-stimulation (left) and CD44 72 hours post-activation (right) of PKM2$^{f/f}$ or PKM2$^{KO}$ T cells. Graph shows mean ± SD, each dot represents one mouse (unpaired t test, n.s. = not significant).
e) Representative Seahorse mitochondrial stress assay of PKM2$^{f/f}$ and PKM2$^{KO}$ T cells at the indicated time points following stimulation with αCD3/αCD28+rIL-2.
f) GFP expression of OT-I transduced PKM2$^{f/f}$ or PKM2$^{KO}$ CD8$^{+}$ T cells (used in **Fig. 5c**).

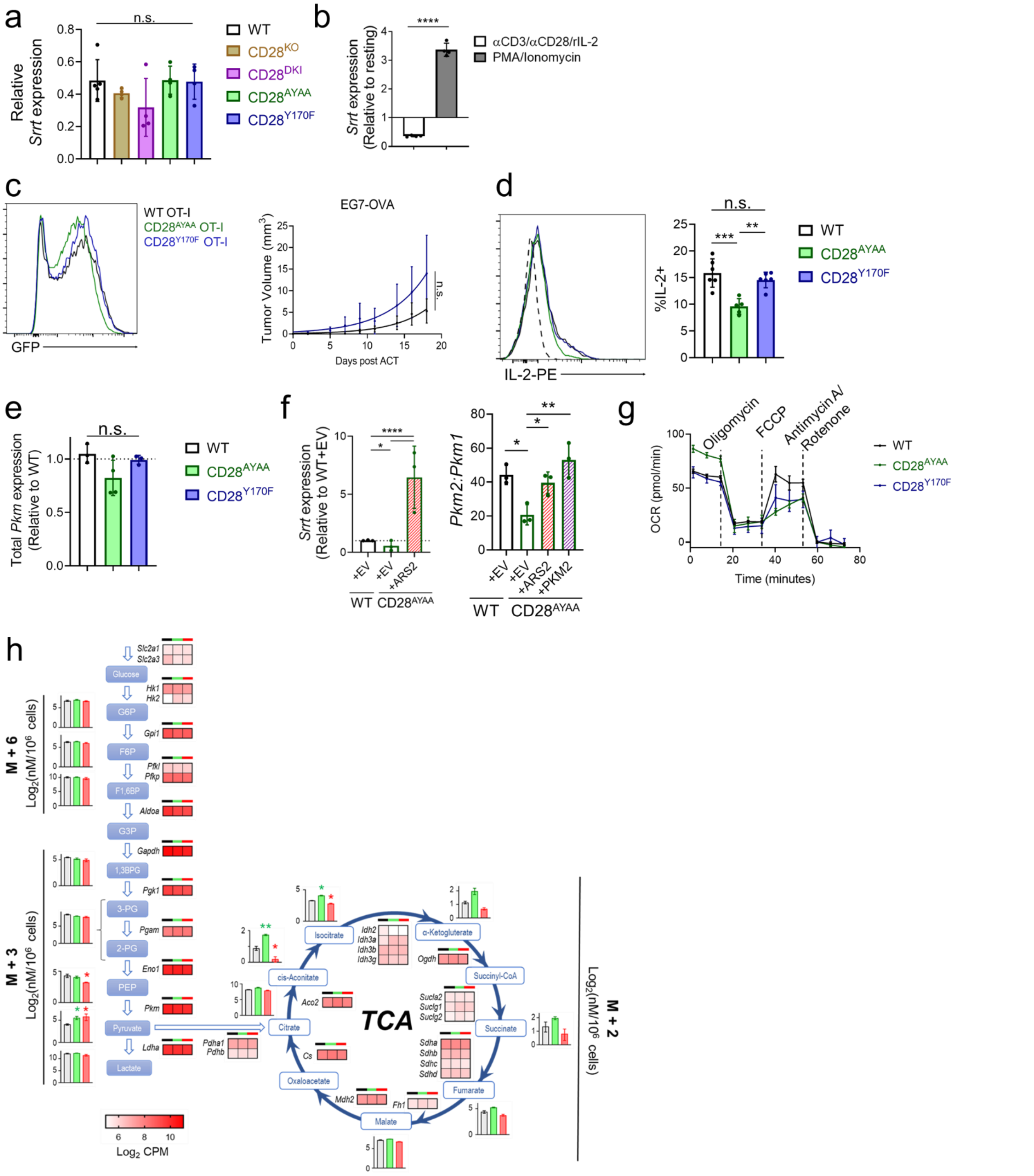

**Supplementary Figure 4:**

a) Basal expression of ARS2 (*Srrt*) in WT and CD28 mutant T cells. Graphs show mean ± SD, each dot represents one mouse (one-way anova, n.s. = not significant).
b) Expression of ARS2 (*Srrt*) in CD28$^{KO}$ T cells stimulated with either αCD3/αCD28 + rIL-2 or PMA/Ionomycin. Graphs show mean ± SD (unpaired t test, ****p<0.0001).

c) Left: OT-I transduction efficiency of WT and CD28 mutant $CD8^+$ T cells as measured by expression of GFP reporter (used in **Fig. 6c**). Right: E.G7-OVA tumor growth of mice receiving WT and $CD28^{Y170F}$ OT-I T cells (excluding $CD28^{AYAA}$ recipients). Graphs show best fit lines and mean tumor volume measurements ± SEM (two-way ANOVA, n.s. = not significant).

d) Flow cytometry assessment of IL-2 expression in activated CD28 mutant effector T cells ($CD8^+CD44^+CD62L^-$) following restimulation with PMA/Ionomycin for 4 hours. Left: Representative flow histogram, dashed line represents FMO. Right: Quantification. Dots represent biological replicates. One-way ANOVA, n.s.=not significant, *p<0.5.

e) Representative graph of total *Pkm* expression in CD28 mutant T cells at 72 hours post-activation (related to **Fig. 6g**). Graphs show mean ± SD of technical replicates, representative of at least 3 independent experiments (one-way anova, n.s. = not significant, ***p<0.001).

f) Left: expression of ARS2 (*Srrt*) in WT or $CD28^{AYAA}$ T cells transduced with empty vector or $CD28^{AYAA}$ T cells transduced with an ARS2 transgene (used in **Fig. 6h**). Right: Ratio of *Pkm2*:*Pkm1* in WT or $CD28^{AYAA}$ T cells transduced with empty vector (EV) or $CD28^{AYAA}$ T cells transduced with retroviral ARS2 or PKM2 (used in **Fig 7d-f**). Graphs show mean ± SD, dots represent individual mice (one-way anova, *p<0.05, ****p<0.0001).

g) Representative Seahorse mitochondrial stress test of WT and CD28 mutant T cells at 72 hours post activation.

h) [U-$^{13}$C] glucose enrichment in glycolytic intermediates and citric acid cycle (TCA) metabolites in WT (black), $ARS2^{KO}$ (red), or $CD28^{AYAA}$ T cells activated for 3 days with αCD3/αCD28 + rIL-2. Bar graphs show mean ± SD of indicated isotopomers of each metabolite measured in 3 biological replicates. Unpaired t tests, *p < 0.05. Heatmaps show expression of indicated genes coding metabolic enzymes.
